# Developmental priming increases copper-tolerance in a model fish species via epigenetic-and microbiome-mediated mechanisms

**DOI:** 10.1101/2025.11.28.691212

**Authors:** Tamsyn M. Uren Webster, Lauren V. Laing, Jemima Onime, Hannah Littler, Rob J. McFarling, Josie Paris, Jennifer A. Fitzgerald, Anke Lange, Audrey Farbos, Karen Moore, Matthew D. Hitchings, Ronny van Aerle, Nic R. Bury, Eduarda M. Santos

## Abstract

Pollution is a significant threat to aquatic ecosystems globally and, in order to survive, natural populations depend upon their ability to rapidly develop tolerance to chemical stressors. We examined whether early-life priming enhances life-long copper-tolerance in a model fish species via developmental plasticity. Stickleback (*Gasterosteus aculeatus*) embryos were pre-exposed to a low concentration of copper (10 µg/L) during early development, reared in clean water for nine months alongside a control group, and then exposed to copper (0,10 and 20 µg/L) for 96 h as adults. Priming markedly reduced evidence of copper-toxicity in adult gills at the transcriptional level (including reduced cellular stress response (CSR) and disruption of ion-homeostasis) and increased inducibility of the metal-binding protein, metallothionein. In parallel, we identified epigenetic and microbiome-mediated mechanisms likely contributing to this tolerance. Pre-exposure induced persistent DNA methylation changes, consistent with priming of CSR and ion-homeostasis pathways. We identified enhanced copper-tolerance in the gill microbiota of primed fish that likely also contributed to host tolerance. These findings provide critical evidence for developmental plasticity induced by chemical stressors in animals, highlight the importance of integrated microbiome and epigenetic responses, and enhance our understanding of how natural populations cope with pollution in their environment.

## Introduction

Aquatic ecosystems are threatened by unprecedented anthropogenic challenges, including chemical pollution. To better understand, predict, and ultimately to mitigate, the impacts of pollution, it is important to consider the relative sensitivity of different species and populations, including their ability to develop tolerance following chemical exposure. Although there is some evidence of local adaptation to pollution in microorganisms, plants and metazoa following long-term exposure^1–4^, for many species survival will ultimately depend on an ability to rapidly acquire tolerance to acute and fluctuating stressors^5,6^.

Phenotypic plasticity allows organisms to rapidly adjust to environmental challenges within their lifetime and encompasses a range of physiological, morphological, and behavioural adjustments^5,7,8^. Acclimation is usually rapidly induced and reversible, with changes diminishing after stressor-removal^5,9,10^. In contrast, developmental plasticity occurs when environmental conditions experienced during critical early-development windows induce persistent, often irreversible, changes in phenotype^10–12^. For plants, there is good evidence that pre-exposure to chemicals and other abiotic stressors during early life, or ‘priming’, can induce persistent tolerance via stress memory^13–15^. For animals, while temperature-induced developmental plasticity has been documented in fish and other ectotherms^7,10,16,17^, it is unclear whether this phenomenon can similarly enhance tolerance to environmental pollutants. Only a handful of studies have examined whether chemical exposure in early development alters subsequent sensitivity^18–21^. This knowledge is critical to understand the sensitivity of natural populations experiencing fluctuating levels of pollution, a common feature of many natural environments.

Epigenetic mechanisms, regulating differences in gene expression, have been shown to underly developmental plasticity across diverse systems^8,22–25^. In plants, multiple epigenetic mechanisms, including chromatin remodelling, DNA methylation and ncRNAs, are known to facilitate primed stress memories, conferring tolerance to environmental stressors, including toxic metals^26,27^. Similarly, for animals, epigenetic mechanisms contribute to increased thermal tolerance following early life exposure^17,25,28,29^. Regarding chemical stressors in animals, research has so far focused only on epigenetic toxicity. Many classes of chemical pollutants, including metals, pesticides and endocrine disruptors, are known to induce epigenetic modifications associated with adverse health outcomes, dependent on the timing and nature of exposure^30–33^. However, the potential for epigenetic mechanisms to contribute to enhanced chemical tolerance in animals remains unexplored.

Host-associated microbiota also play a critical role in influencing sensitivity to environmental stressors by extending host adaptive phenotypic capacity^34,35^. While microbiomes are often sensitive to disruption by environmental stressors, they have an extensive capacity to rapidly develop tolerance^34,36,37^. Crucially, microbial adaptive plasticity can enhance host tolerance to environmental challenges^38,39^. Tolerant microbiota can, for example, limit adverse effects in the host associated with microbiome disruption, and/or confer specific benefits, such as enhanced metabolism or sequestration of toxins^36,40,41^. As with the epigenome, the microbiome has heightened environmental sensitivity during early development^42^. While microbiome priming is emerging as a technique to enhance agricultural productivity and stressor resilience^41^, its potential role in environmental chemical-tolerance is largely unexplored.

It is unknown whether, in animals, tolerance to environmental chemicals can be acquired via developmental plasticity, the extent to which this occurs, or the specific mechanisms contributing to this effect. We aimed to address these questions by examining whether copper, a widespread aquatic pollutant, can induce developmental plasticity in three-spined stickleback *(Gasterosteus aculeatus*), a well-established model in evolutionary ecology and ecotoxicology. We tested the hypothesis that priming would induce persistent, elevated tolerance to copper, with both epigenetic and microbiome-mediated mechanisms contributing to this adaptive response. To specifically examine the capacity for developmental plasticity, distinct from acclimation, we exposed stickleback embryos to an environmentally-relevant concentration of copper during early development, returned them to control conditions for nine months (until maturity), and then compared response to copper exposure in primed and naïve adults.

## Results

### Early-life exposure promotes continued copper accumulation in the gill

Stickleback embryos were pre-exposed to an environmentally relevant concentration of copper (*nominal:* 10 µg/L, *measured* 11.4 ±0.3 µg/L) during early development (one-cell stage to hatched larvae; 1-217 hpf), alongside a synthetic freshwater control group (*measured* 0.2 ±0.004 µg/L Cu). Survival was high in both groups, but copper exposure caused a small increase in embryo/larval mortality rate (Naïve: 0.69%, Pre-exposed: 1.24%; P=0.0384). Pre-exposure also increased larval whole-body copper concentration (t =-9.20, df = 5.58, P<0.001; Figure 1a).

**Figure 1.**
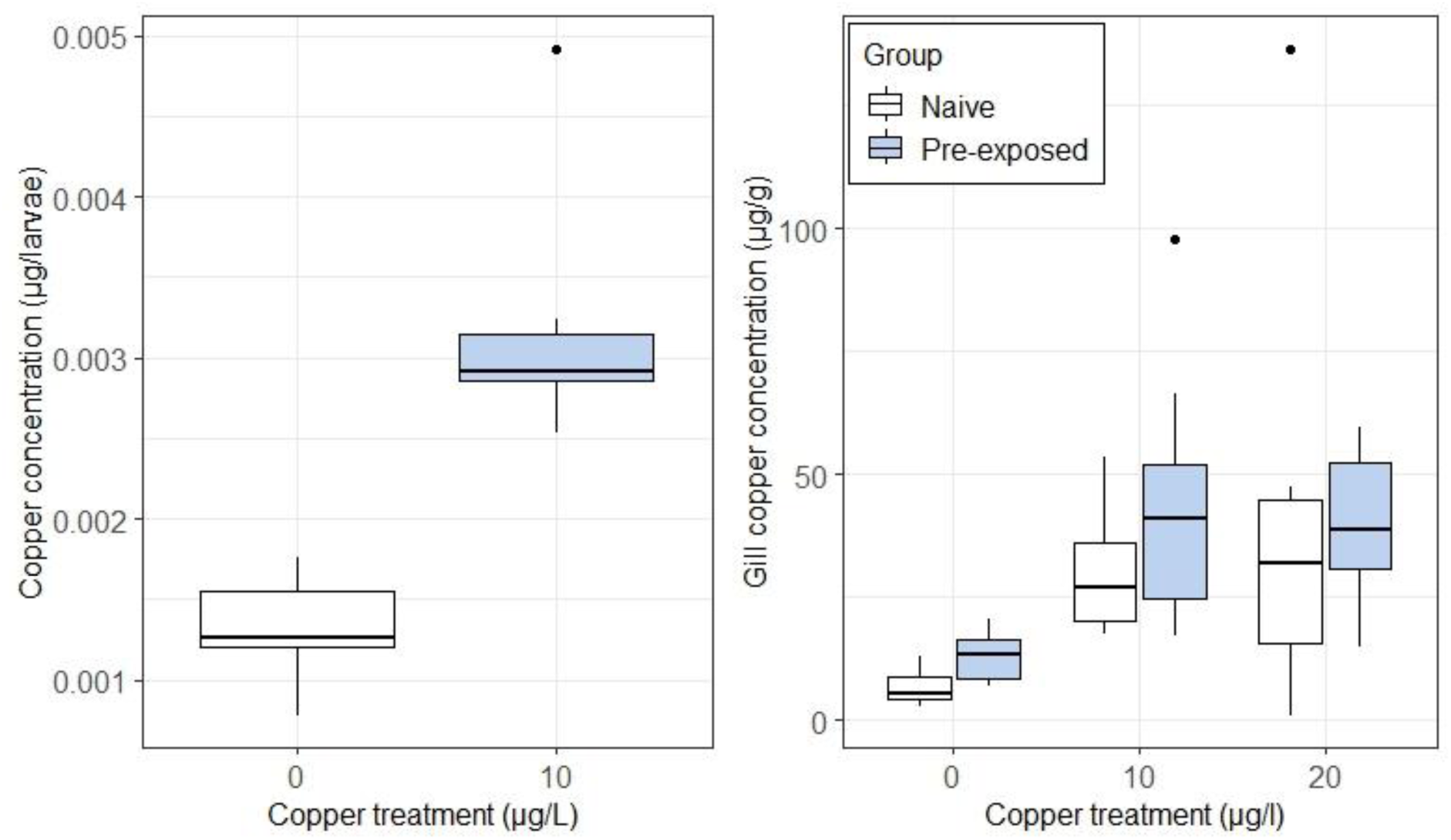
Copper measured in. **A)** whole larvae after copper exposure during embryonic development (n= 12 pools of 5 larvae/group) and **B)** in the gills of stickleback from each group later exposed to copper for 96-h as adults after 9 months depuration in clean water (n= 10/group).

After nine months depuration in clean water, both naïve and primed adults were (re-)exposed to two concentrations of copper (*nominal:* 10 µg/L, *measured:* 13.5 ±0.3 µg/L; and *nominal:* 20 µg/L, *measured:* 21.8 ±0.05 µg/L) for 96-h alongside a control (*measured:* 5.0 ±0.05 µg/L). Pre-exposed adult stickleback had accumulated higher concentrations of copper in their gills compared to the naïve fish, while adult exposure also increased gill copper concentration in both groups (*Pre-exposure:* F_1,48_=8.55, P=0.005, *Adult-exposure:* F_2,48_=21.18, P<0.001, *Interaction:* F_2,48_ =0.36, P=0.69; Figure 1b). In contrast, there was no discernible effect of either pre-exposure or adult exposure on the concentration of copper measured in muscle or liver tissue. No mortalities or behavioural changes were observed during the adult copper exposure, and neither pre-exposure nor adult exposure to copper affected fish size.

### Pre-exposure substantially reduces and modifies transcriptional stress response to copper

We focused the molecular analyses on the gills of adult fish, given their role in metal uptake and the measured differential accumulation of copper in this tissue. We conducted transcriptomic profiling in both the naïve and primed groups following (re-)exposure to 0 and 10 µg/L copper.

Pre-exposure to copper during embryonic development had minimal lasting effects on baseline transcription in adult fish, with only two differentially expressed genes (DEGs) identified between primed and naïve fish. Gene Set Enrichment Analysis (GSEA), identified a limited number (seven) of negatively enriched GO terms (Table S2); which were all related to cytoskeleton structure and function (actin, myosin, troponin, calcium binding).

We then compared the transcriptomic response to copper exposure of primed (pre-exposed) adults with that of naïve fish (adult fish exposed to copper for the first time). The magnitude of transcriptional response was far greater in naïve fish than in primed fish (1575 and 45 DEGs, respectively; Figure 2, Table S1). Of these, 18 DEGs were common between groups, including those encoding seven heat shock proteins (HSPs; subtypes 90,70 and 30). These molecular chaperones, critical in cellular stress response, were the most significantly up-regulated genes in response to copper in both groups, but the magnitude of this up-regulation was markedly higher in naïve fish (ranging 7-805 fold increase) than in pre-exposed fish (ranging 3-170 fold increase). Metallothionein B, a metal-sequestering protein, was also strongly up-regulated in both groups, although, in this case, by a greater magnitude in the pre-exposed fish (5.9 fold increase) compared to naïve fish (3.3 fold increase).

**Figure 2.**
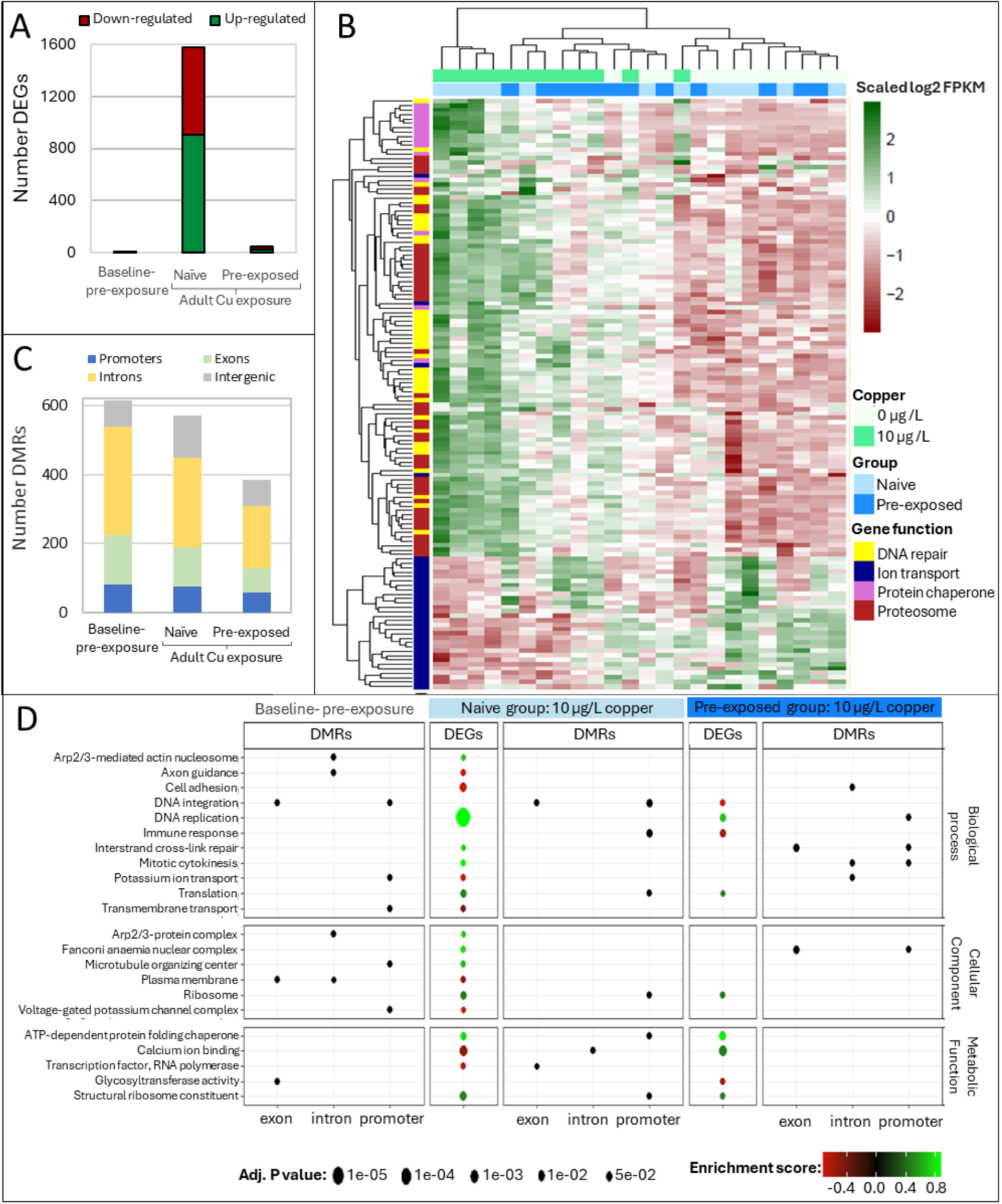
**A**) Number of significantly differentially expressed genes (DEGs) identified in response to pre-exposure alone (baseline) and in naïve or pre-exposed groups exposed to 10 µg/L copper as adults. **B)** Heatmap visualising the expression of selected DEGs, based on their primary function (N= 135 DEGs out of a total of 1608 identified in response to copper exposure across both groups (see Fig S1). **C)** Number and the genomic context of differentially methylated regions (DMRs) identified in response to pre-exposure alone (baseline) and in both naïve and pre-exposed groups exposed to 10 µg/L copper. **D)** Shared enriched GO terms associated with both DMRs and DEGs.

GSEA revealed a greater magnitude of response to copper in naïve fish (63 enriched terms) than in pre-exposed fish (34 terms). In naïve fish there was strong enrichment of ‘DNA replication’, ‘Cell-cycle’ and associated terms, as well as terms related to protein refolding and synthesis. Similar terms were enriched in pre-exposed fish, but to a far lesser extent. Processes regulated exclusively in naïve fish included strong enrichment of those associated with DNA repair and the proteosome, while terms associated with ion homeostasis were supressed, reflecting down-regulation of >30 genes encoding potassium, sodium, calcium, magnesium, ammonium and bicarbonate channels and cotransporters. A further marked distinction between the response of each group was that terms associated with cytoskeleton (including actin, myosin, troponin and calcium ion binding) and extracellular matrix (ECM) interactions were supressed in naïve fish but enhanced in pre-exposed fish.

### Copper exposure induced marked and long-lasting changes in the gill methylome

We conducted genome-wide DNA methylation profiling (RRBS) in the gills of naïve and primed fish after (re-)exposure, on the same samples used for transcriptomic analysis. In contrast to that observed for transcription, we measured considerable, lasting changes in the gill methylome of adult fish following developmental pre-exposure to copper and in the absence of any subsequent exposures. A total of 615 differentially methylated regions (DMRs) were identified between the pre-exposed and naïve groups (328 hyper-methylated, 287 hypo-methylated). Of these, 13.5 % overlapped putative promoters (pps), while 20.4% and 42.2% were associated with exons and introns, respectively (Figure 2c, Table S3). Among the genes associated with these DMRs (overlapping pps, exons, introns), notable examples included those involved in ion homeostasis (particularly sodium, potassium and calcium transport), metal transport and binding (including those encoding copper-uptake protein 2, ceruloplasmin and ferritin), those with immune function, and a number of lncRNAs. GSEA, performed separately for different genomic contexts, identified 41, 50 and 45 enriched terms associated with DMRs located within pps, exons and introns, respectively (Table S4, Figure S3). Among the most enriched terms were those associated with membrane transport and ion homeostasis, including copper-ion transport. Regulation of immune response (interleukin production) and many terms associated with cellular growth and division, cell adhesion and signalling, were also evident.

Adult copper exposure also induced considerable changes in the methylome, and these were more extensive in naïve fish. A total of 571 DMRs (287 hyper-methylated, 284 hypo-methylated) were identified in naïve fish exposed to copper, compared to 385 DMRs (202 hyper-methylated, 183 hypo-methylated) in pre-exposed fish (Figure 3, Table S3). Functional enrichment analysis identified a greater number of terms associated with DMRs in the naïve fish (Table S4, Figure S3). These included, most strongly, ‘immune system’, several terms related to chaperone-mediated protein refolding and, more broadly, many terms related to protein, nucleic acid and cellular repair and turnover. In pre-exposed fish, there were some broad similarities in the function of enriched terms to those in the naïve fish, including those related to nucleosome, immune response and cellular turnover but, notably, ‘ion transport’ was only enriched in pre-exposed fish.

**Figure 3.**
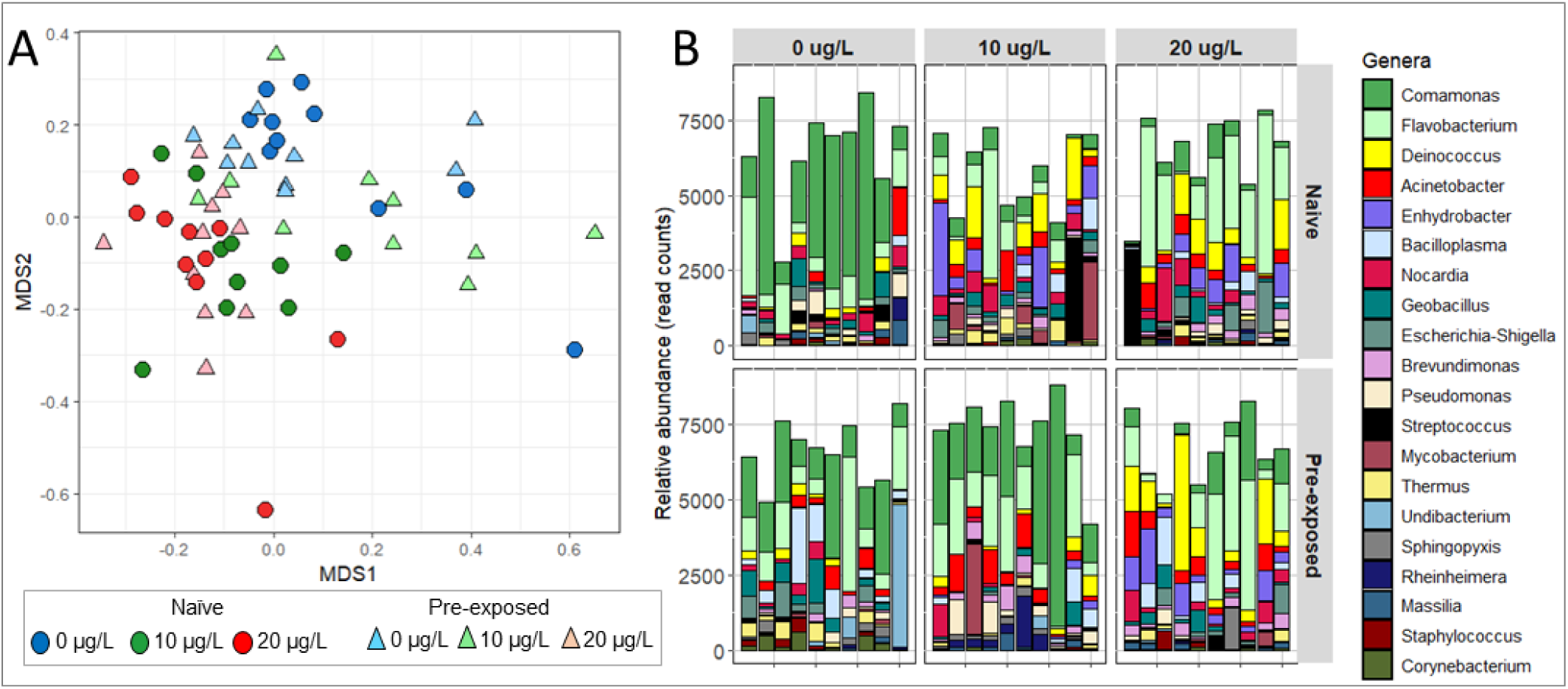
**A**) Gill microbial community structure in adult sticklebacks (re-)exposed to 0, 10 or 20 ug/L copper, visualised using Bray-Curtis dissimilarity values, and **B)** Relative abundance of the top 20 most abundant bacterial genera identified across all samples.

### Genes & functional pathways with both epigenetic and transcriptional modifications

DMRs were identified within the gene body or putative promoter of 13 DEGs in naïve fish exposed to copper as adults, and one DEG in pre-exposed fish exposed to copper (Table S5). Functions of these genes included protein degradation, synthesis, folding and damage-repair, as well as calcium signalling, cytoskeleton, and the regulation of cell cycle and cell movement. There were also six shared enriched GO terms associated with both altered methylation and transcription in naïve fish (relating to protein synthesis, folding and calcium signalling) and one shared term (DNA replication) in pre-exposed fish (Figure 2D).

We hypothesised that persistent methylation differences following pre-exposure influenced the transcriptional responses of adult stickleback (re-)exposed to copper. We identified 13 genes with differential baseline methylation following pre-exposure that showed a different transcriptional response to copper between primed and naïve adult fish; all of these genes were only transcriptionally responsive to copper in the naïve fish (Table S5). The functions of these genes were related to protein degradation and synthesis, DNA repair, regulation of cell cycle and cell movement, as well as cytoskeleton and regulation of ion channels. Eight shared enriched GO terms, related to cytoskeleton and potassium ion transport, were also identified (Figure 2d).

### Early-life priming increases gill microbiota copper-tolerance

We hypothesised that priming during the early stages of microbiome establishment would promote enrichment of gill-associated microbiota better able to withstand copper exposure. To test this, we characterised the gill microbiomes of primed and naïve adult fish exposed to 0, 10 and 20 µg/L. Exposure to the higher concentration of copper disrupted microbiome community structure (Bray-Curtis dissimilarity) in all fish, regardless of priming, but the effects of the lower copper concentration differed between groups (*Adult exposure:* F_1,56_ =5.938, P <0.001, *Pre-exposure:* F_1,56_ =1.221, P=0.189, *Interaction:* F_1,56_ =1.55, P=0.0493; Figure 3). While fish from the naïve group exposed to 10 µg/L showed microbiome disruption similar to those exposed to 20 µg/L copper, the microbiomes of primed fish were more resistant to change and remained similar to those of the fish unexposed to copper as adults.

We examined the differences in community composition contributing to these structural changes. We identified two amplicon sequence variants (ASVs) with baseline differential abundance between naïve and pre-exposed fish (*Vibriomonas* and *Candidatus Bacilloplasma*).

In adults exposed to copper, pre-exposure reduced the number of differentially abundant ASVs identified (14 and 29 following exposure to 10 and 20 µg/L, respectively; Table S6) compared to in naïve fish (32 and 40 following exposure to 10 µg/L and 20 µg/L; Table S6). Notably, the most abundant ASV overall, *Comamonas* sp., was strongly inhibited by copper in naïve fish (reduced by 13– and 62-fold following exposure to 10 and 20 µg/L), but not in pre-exposed fish (2-fold reduction in response to 20 µg/L only). At the same time, there was a marked increase in the abundance of ASVs from the genera *Deinococcus, Enhydrobacter, Acinetobacter, Flavobacterium* and *Brevundimonas* in both groups, but generally the magnitude of increase was higher in naïve fish.

There were no detectable effects of pre-exposure or adult exposure on overall richness or diversity of ASVs present (Chao1 richness-*Pre-exposure:* F_1,56_=2.05, P=0.158, *Adult-exposure:* F_1,56_=0.49, P=0.485; Shannon diversity-*Pre-exposure:* F_1,56_=0.02, P=0.880, *Adult exposure:* F_1,56_=0.09, P=0.768, *Interaction:* F_1,56_=1.00, P=0.32).

## Discussion

We examined whether early-life priming induced developmental plasticity in stickleback using copper as a model toxicant, hypothesising that epigenetic and microbiome-mediated mechanisms contribute to this phenomenon. We found that priming substantially ameliorated copper toxicity in adult fish, an effect characterised by reduced stress response at the transcriptional level. In parallel, pre-exposure induced considerable, and persistent, changes in the gill methylome, with notable similarities between differentially methylated and expressed gene pathways suggesting a priming effect on cellular stress response pathways. Furthermore, early-life priming markedly increased copper-tolerance in the gill microbiome, likely also reducing the toxic effects of copper exposure on the stickleback host.

### Early-life priming increased tolerance to copper toxicity in adult fish

Transcriptional response to copper was markedly reduced in adult fish that had been primed in early life. In both naïve and pre-exposed fish, we identified transcriptional changes dominated by genes and pathways associated with the cellular stress response (CSR), but the magnitude of transcriptional changes and enrichment scores were far greater in naïve fish. The CSR is highly and broadly inducible by many stressors, but its nature varies depending on the severity and duration of the stressor^43,44^. In both groups, although with a greater magnitude in naïve fish, we characterised an extensive compensatory CSR, reflecting the repair of cellular components. As part of this, we identified a marked increase in the transcription of heat shock proteins and other molecular chaperones responsible for refolding of damaged proteins, as well as genes involved in DNA repair pathways. There was also strong up-regulation of DNA replication, protein synthesis and cell division pathways, consistent with an increase in cellular turnover. In naïve fish only, there was a distinct enrichment of the proteosome, responsible for degradation of irreversibly damaged proteins. This supports the induction of a more severe CSR in naïve fish, characterised by a switch from repair to autophagic pathways, indicating that greater copper-induced cellular damage occurred in these fish.

Further evidence that priming reduced copper toxicity included markedly different transcriptional effects associated with ion transport and cytoskeleton dynamics. In naïve fish only, we found down-regulation of a suite of ion transporters, particularly potassium channels, indicating broadscale disruption of ion homeostasis, a well-known mechanism of copper toxicity, especially in the gills^45^. Metals can induce cytoskeleton toxicity via interference with calcium signalling and its regulation of troponin-tropomyosin-actin dynamics^46–48^. Consistent with this, in naïve fish only, we found inhibition of these pathways, indicative of substantive toxicity. However, in pre-exposed fish, there was instead a consistent stimulatory effect, likely associated with a compensatory CSR, encompassing transcripts associated with enhanced cellular signalling, reorganisation and stability^43,44^.

In contrast to the reduced magnitude of CSR, we found that metallothionein, a key metal-binding protein that sequesters and reduces the toxicity of free metal ions, was more strongly up-regulated in pre-exposed fish in response to copper exposure. A similar priming effect, increasing inducibility of this protein, has been previously associated with enhanced tolerance to toxic metals in populations with different exposure histories^49^ and, more widely, the increased inducibility of genes with protective functions contribute to developmental plasticity and increased stressor tolerance^50^. Importantly, the minimal differences in baseline transcription identified between the naïve and pre-exposed fish are consistent with persistent developmental plasticity, rather than acclimatory effects (i.e. frontloading transcription)^50^.

Importantly, we found no effects on long-term survival or growth in primed fish compared to their naïve counterparts, suggesting that developmental plasticity in copper tolerance was not associated with overt energetic costs. Surprisingly, early-life exposure caused fish to continue to accumulate more copper in their gills during the nine-month depuration phase, indicative of changes in metal-homeostasis physiology. Following pre-exposure, we identified persistent methylation differences in *slc31a2* (copper-uptake protein), ceruloplasmin (copper-transport protein) as well > 20 sodium and calcium transporters and channels, which are also responsible for a substantial amount of copper uptake in fish gills^51^, although there were no differences in their baseline transcription. Increased copper accumulation was only evident in the gills suggesting that primed fish may have an increased tendency to sequester copper in this tissue, likely in a less-toxic and/or bioavailable form, and consistent with the higher inducibility of metallothionein proteins identified.

### Epigenetic mechanisms contribute to developmental plasticity in copper-tolerance

Pre-exposure to copper caused extensive and persistent changes in the gill methylome, evident even after nine months in control conditions. This has not been reported before for metals but is consistent with existing evidence that developmental thermal stress induces long-lasting changes in DNA methylation in fish^25,52^, and, more broadly, with there being heightened epigenetic sensitivity in early life which, when combined with environmental fluctuation, drives phenotypic plasticity^24^. There were also extensive changes in the adult gill methylome immediately following copper exposure, including within many of the same gene pathways modulated by pre-exposure. This suggests that DNA methylation plays a similar role in acute stress responses as in developmental priming, an effect previously identified in stickleback challenged with thermal stress^25^.

We identified a considerable functional overlap in epigenomic and transcriptomic responses to copper, including in many gene pathways broadly associated with the CSR (especially protein turnover, cytoskeleton regulation and cell cycle) and ion transport. Thirteen genes were both differentially methylated and expressed in acute response to copper exposure in adults, while another 13 genes, that were persistently differentially methylated following pre-exposure, subsequently showed different transcriptional responses to copper in primed and naïve fish. In addition, we identified similarities in modulated gene pathways, which were predominantly associated with the CSR and ion homeostasis. Together, our results suggest that epigenetic regulation has a priming effect on CSR and ion homeostasis pathways and subsequently contributes to the reduction in transcriptional stress response identified in pre-exposed fish. Priming could, for example, facilitate a more efficient and/or readily inducible CSR, similar to that which occurs in plants, where developmental stress priming induces diverse methylation changes within CSR, defence and signalling pathways that are associated with improved tolerance to many stressors, including metals^53^.

### Priming increased copper tolerance of the gill microbiota

Copper exposure caused a dose-dependent, disruptive effect on gill microbial community structure, that was more severe in naïve fish. Early-life priming reduced the number of differentially abundant ASVs identified in response to both concentrations of copper and, in particular, protected against significant community disruption by the lower (10 µg/L) concentration. The disruptive effects of copper were characterised by a substantial decline in the presence of the otherwise most abundant community member, *Comamonas* sp. In parallel, there was an increased prevalence of genera including *Deinococcus,* known for its high abiotic stressor tolerance^54^, as well as *Acinetobacter* and *Flavobacterium,* both of which are genera that include opportunistic fish pathogens and have previously been associated with microbiome stressor-disruption^42,55^. Our results support the hypothesis that priming enabled commensal members of the core microbiome, including *Comamonas,* to develop enhanced copper tolerance, likely through mechanisms such as genetic adaptation, gene transfer or plasticity^41^. This may have been further reinforced by elevated copper accumulation occurring in the gills of primed fish, providing a sustained selective environment for tolerant strains.

We propose that the extensive changes in community structure observed were associated with disrupted gill microbiome function, thereby increasing the toxic effects of copper to the host. Considering the increase in opportunistic pathogens observed, adverse effects on host physiology could include a disruption of normal microbial contribution to pathogen defence and provision of beneficial metabolites, and exacerbation of host inflammatory stress responses. Having a more tolerant, less-disrupted microbiome, is likely to be beneficial to the host. In the context of the holobiont concept, our results support the hypothesis that microbiome tolerance also contributes to increased stickleback copper-tolerance. Further research should establish whether microbiota additionally provide specific copper-adaptive benefits to the host, such as sequestration of metals, as reported in plants^41^.

### Conclusions

Developmental plasticity, together with acclimation, transgenerational plasticity and genetic adaptation, contribute to variations in the sensitivity of natural populations to environmental stressors, ultimately influencing their ability to survive^50^. Establishing the capacity for organisms to acquire tolerance, and elucidating underlying molecular mechanisms, is therefore essential to understanding and predicting the risks posed by pollution and other stressors in the natural environment. Here, we provide some of the first evidence for developmental plasticity in chemical tolerance in animals and demonstrate that both epigenetic and microbiome-mediated mechanisms are associated with this effect.

## Methods Summary

### Ethics approval

All experiments were approved by the University of Exeter Ethics committee and conducted under licence from the UK Home Office according to ASPA.

### Embryo copper experiment

Pools of 50 embryos were exposed to either a water control (0 µg/L copper) or 10 µg/L copper (added as CuSO_4_) from 1-217 hours post fertilisation (covering the period of embryogenesis and hatching, and including the period of epigenetic reprogramming and microbiome colonisation). Exposures were conducted in 500 ml acid-washed glass dishes containing aerated synthetic freshwater^56^, with four replicates per treatment, repeated three times with different parental fish (see SI). Embryo mortalities and hatching were recorded daily. Water samples were collected at 120 and 217h, and 20 larvae from each replicate (n=12) were collected at 217h for copper measurement (see SI). Embryo survival and copper uptake were analysed using a student’s t-test in R (v4.3.3; ^57^). 200 larvae from each of the control and copper-exposed groups were then maintained in dechlorinated tap water (in duplicate tanks per group) for nine months.

### Adult copper exposure experiments

Adult male stickleback from both the naïve and pre-exposed groups were exposed to either 0, 10 or 20 µg/L copper for 96 hours. Each treatment was performed in duplicate 40 L tanks, with nine fish per tank, supplied with flow-through dechlorinated tap water. Fish were not fed for the duration of the exposure. Water samples were collected at 24 and 72h for copper measurements (see SI). After exposure all fish were humanely sacrificed by lethal dose of benzocaine (0.5 g/L; Sigma-Aldrich), followed by destruction of the brain. Gill, liver and muscle tissue were snap frozen and stored at –80°C. All left-side gill arches were used for copper-content analysis (see SI), while all right-side gill arches were used for molecular analysis. The effect of both pre-exposure and adult exposure on fish weight and tissue copper concentration was examined using ANOVA in R.

### Sequencing & Bioinformatics

Transcriptome, methylome and microbiome analyses were conducted on gill tissue from adult fish, with full details in SI. Briefly, for transcriptome and methylome analyses, gill RNA and DNA were co-extracted using Qiagen AllPrep DNA/RNA Mini kits from fish from four treatment groups (naïve and pre-exposed fish exposed to 0 and 10 µg/L Cu; n=6 per group). RNA-seq libraries were prepared using an Illumina TruSeq Stranded RNA Sample Preparation kit and sequenced using an Illumina HiSeq 2500 (100 bp paired end). RRBS libraries were prepared using Ovation RRBS Methyl-Seq kit (Tecan Systems) and sequenced using an Illumina NovaSeq (100 bp paired end). For microbiome analysis, DNA was extracted from gill tissues using the Qiagen PowerSoil DNA

Isolation Kit (n= 10 from all six treatment groups). Libraries were prepared, amplifying the 16S rRNA V4 region using primers 515F, 806R ^58^, based on the Illumina 16S Sequencing Library Preparation protocol^59^ and sequenced using an Illumina MiSeq (300 bp paired end).

RNA-seq reads were quality filtered using Fastp (v0.23.1.3;^60^) and aligned to the *Gasterosteus aculeatus* reference genome^61^ using STAR (v2.7.9a;^62^). Mapped reads were quantified with RSEM (v1.3.1;^63^) and differential gene expression analysis was performed using DESeq2 (v1.38.3;^64^). Gene set enrichment analysis (GSEA) was conducted using ClusterProfiler (v4.7.1;^65^), incorporating customised GO term annotations generated using InterProScan (v5.55.88.0;^66^) and Blast2Go (v1.4.12;^67^).

RRBS reads were quality filtered using TrimGalore^68^ and aligned to the reference genome with Bismark (v0.23.1;^69^. Differentially-methylated regions (DMRs) were identified using DSS (v2.48.0;^70^). Genomic location of DMRs (classified as putative promoters (within 1000 bp of the transcription start site), exons, introns, or intergenic regions), and gene annotations were determined using Genomation (v3.17;^71^). GSEA was performed using g:Profiler^72^ using input gene lists ranked by methylation fold change.

16S rDNA data were processed using DADA2 ^73^ within Qiime2 (v2024.2,^74^). Reads were quality-filtered, merged, de-noised, assigned to amplicon sequence variants (ASVs) and taxonomically classified using the Silva reference database (v132;^75^). Alpha and beta diversity metrics were calculated using the Vegan package^76^. The effects of both early-life priming and adult copper exposure on Chao1 richness and Shannon diversity were assessed using ANOVA. Bray-Curtis dissimilarity was analysed using PERMANOVA and differential ASV abundance was evaluated using DESeq2 ^64^.

## Author contributions

LL, TUW & ES conceived & designed the study. LL conducted the stickleback experiments with help from JF, JP and AL. LL, HL, RMF, AL & TUW conducted molecular work and AF, KM and MH conducted the Illumina sequencing. NB and LL conducted the metal analysis. TUW (RRBS & 16S), JO (RNA-seq) and LL (metal content) led the data analyses with contributions from RvA, JP, NB & ES. ES supervised the study and, together with LL & TUW, led funding acquisition. TUW wrote the original draft. All authors contributed to review and editing of the final manuscript.

## Acknowledgements

We thank staff from the University of Exeter Aquatic Resources Centre for assistance with fish husbandry. Funding was received from the Fisheries Society of the British Isles (FSBI), the BBSRC (BB/S004300/1), NERC PhD studentships, the Exeter-Cefas strategic alliance, Cefas Seedcorn, and the Swansea University College of Science Research Fund. RNA-seq and RRBS sequencing was performed at the University of Exeter Sequencing Service, which utilised equipment funded by the Wellcome Trust Institutional Strategic Support Fund (WT097835MF), Wellcome Trust Multi-user Equipment Award (WT101650MA) and BBSRC LOLA award (BB/K003240/1).

