## Supplementary material for "Developmental priming increases copper-tolerance in a model fish species via epigenetic-and microbiome-mediated mechanisms": SI methods

***Supporting Information- extended material and methods***

*Fish maintenance*

Prior to the experiment, adult three-spined stickleback, originating from the River Erme, UK, were kept in mixed sex groups (20 fish per 112 L tank), supplied with aerated synthetic freshwater at 15 ± 1°C. Fish were fed twice daily with blood worm (*Chironomus* sp.), and once daily with live *Artemia nauplii*, to satiation. Pools of stickleback embryos were obtained via *in vitro* fertilisation[1] from 20-26 females and 3-5 males and randomly allocated to treatments to ensure both genetic variability within the experiment and equity across treatment groups. Embryos were incubated in aerated synthetic freshwater (according to the ISO-7346/3 guideline, ISO water, diluted 1:5 [2]). During the priming experiment, daily water changes were performed to maintain oxygen saturation >80%.

After the priming experiment, larval fish were fed twice daily with ZM dry food of appropriate grade, and once daily with live *Artemia nauplii,* to satiation, and then the adult diet described above.

*Metal concentration measurement*

All samples for metal analysis were collected in acid washed tubes. Tank water samples where acidified with nitric acid (70%, ≥99.999% purity, Sigma Aldrich) to a final concentration of 0.01%. Larval (pools of 20, n= 12 per treatment) and adult gill, muscle and liver (n= 8-10 per treatment) samples were freeze dried, the dry mass determined, then digested with 500µl of nitric acid for

48h with vortexing to ensure complete digestion. 0.1% hydrogen peroxide was added to remove fats, and the digested solution was then diluted (1:10) with ultrapure water. Copper concentration in all water, larval and gill samples were measured by ICP-MS using a Perkin Elmer NexION 350D instrument running Syngistix software, v1.0.

*DNA/RNA extraction and library preparation*

RNA and DNA QC was performed using a NanoDrop ND-1000 Spectrophotometer (NanoDrop Technologies, Wilmington USA) and an Agilent 2200 TapeStation (Agilent Technologies, CA, USA). RNA-seq libraries were prepared from 500 ng RNA (RIN ≥8.6) using the Illumina TruSeq Stranded RNA Sample Preparation kit according to the manufacturer's instructions, with ERCC spike-in controls (Ambion, TX, US) added to each library. Libraries were sequenced using an Illumina HiSeq 2500 (100 bp paired end reads). RRBS libraries were prepared using Ovation RRBS Methyl-Seq kit (Tecan Systems) from 100 ng input DNA. 5-Methylcytosine (0 and 100% methylation) DNA standards (Zymo Research) were spiked into the libraries according to the manufacturer's instructions. Libraries were sequenced using an Illumina NovaSeq platform (100 bp paired end reads).

Microbiome library preparation was performed, amplifying the 16S rRNA V4 hypervariable region (primers 515F, 806R [3]), as described in [4] based on the Illumina 16S Metagenomic Sequencing Library Preparation protocol [5] using Platinum Hot Start Taq Polymerase (Invitrogen). All libraries were purified using AMPure XP magnetic beads (Beckman Coulter), indexed using Nextera XT indices (Illumina), multiplexed in equimolar concentrations and sequenced using an Illumina MiSeq platform (300 bp paired end). Two extraction blanks, and two PCR blanks were prepared and sequenced alongside the gill samples.

*RNA seq data analysis*

Paired-end reads were subjected to initial quality trimming, filtering and adaptor removal with Fastp (v0.23.1.3; [6]), with exclusion of reads shorter than 60 bp following 3' window trimming (min Phred score 22). Filtered paired reads were aligned to the *Gasterosteus aculeatus* v.5 reference genome (NCBI RefSeq GCF_016920845.1 [7]) using STAR (v2.7.9a; [8]). Gene quantification of uniquely aligned reads was performed using RSEM (v1.3.1; [9]). Alignment and quantitation parameters were adopted from the NASA GeneLab RNA-seq consensus pipeline [10].

Differential gene expression analysis was performed using DESeq2 (v1.38.3;[11]), including pre-filtering to retain only genes with at least one count per million in a minimum of six samples. Pairwise comparisons (Wald tests) were performed to test the effects of i) early-life exposure to copper (Naïve-0 v Pre-exposed-0); ii) adult copper exposure in naïve fish (Naïve-0 v Naïve-0.01), and iii) adult copper exposure in pre-exposed fish (Pre-0 v Pre-0.01). Genes were considered differentially expressed if they had an adjusted p-value < 0.05 (Wald test). Gene set enrichment analysis (GSEA) was conducted using ClusterProfiler (v4.7.1; [12]). Significantly enriched Gene Ontology (GO) terms were identified using an adjusted p-value threshold of < 0.05 (Benjamini–Hochberg correction). Custom GO term annotation for the *G. aculeatus* v.5 reference genome [7] was generated using InterProScan (v5.55.88.0; [13]) and Blast2Go from the OmicsBox (v1.4.12; [14]) platform.

*RRBS data analysis*

Paired-end reads were initially subjected to quality filtering. Adaptors were removed with TrimGalore [15], followed by trimming and filtering with the NuGEN script, trimRRBSdiversityAdaptCustomers.py (https://github.com/nugentechnologies/NuMetRRBS).. After processing, 940.7 million paired reads remained, ranging from 25.9- 56.6 million reads per sample, and these were aligned to the *G. aculeatus* v.5 reference genome [7] with Bismark (v0.23.1; [16]) in directional mode, with Bowtie2 (v2.45.1;[17]) and Samtools (v1.15.1;[18]). Overall and unique paired end genomic alignment rates were 77.3 ± 0.9% and 63.1 ± 1.36 % (Table S1). Only uniquely aligned paired reads were retained for further analysis.

Differentially-methylated regions (DMRs) were identified using the DSS package in R (v2.48.0; [19]). To match the approach used for the RNA-seq analysis, the effects of i) early-life exposure to copper (Naïve-0 v Pre-exposed-0); ii) adult copper exposure in naïve fish (Naïve-0 v Naïve-0.01) and iii) adult copper exposure in pre-exposed fish (Pre-0 v Pre-0.01) were tested. Wald tests were initially performed to identify differentially methylated CpG sites (DMCpGs), using CpG smoothing and a relaxed threshold ($P<0.05$), with all (non-filtered) data included. Putative DMRs were then predicted based on clustering of DMCpGs. Stringent criteria were used to define DMRs; only those regions containing ≥4 CpG sites, with a minimal length of 50 bp, and exhibiting ≥20% methylation difference between groups were included.

Genomic locations of DMRs, classified as putative promoters (within 1000 bp of the transcription start site (TSS)), exons, introns or intergenic regions, and associated gene annotations were identified using Genomation (v3.17; [20]) and a preformatted BED file downloaded from the UCSC Table Browser [21] https://genome-euro.ucsc.edu/cgi-bin/hgTables ). For DMRs located within putative promoters, exons, or introns, the nearest TSS was identified and used to assign gene IDs. GO term enrichment analysis was performed using g:Profiler [22] with input gene lists ranked by methylation fold change and annotated using the custom GO terms described for the RNA-seq analysis.

*Microbiome data analysis*

16S rDNA sequence data were analysed using DADA2 [23] within Qiime2 (v2024.2, [24]). Primer sequences were removed, and reads were 3' truncated to 280 bp (forward) or 240 bp (reverse) based on quality scores. Reads were merged, de-noised, screened for chimeras, and assigned to amplicon sequence variants (ASVs). Taxonomic classification of ASVs was performed using the SILVA reference database (v132; [25]) using a custom trained classifier [26]. Eukaryotic and chloroplast sequences were removed, resulting in a total of 1,719 unique ASVs. All samples were subsampled to an equal depth of 6,946 high-quality reads prior to downstream analysis using the Vegan package [27].

The effects of early-life (pre-exposure) and adult copper exposure on microbiome alpha diversity (Chao1 richness and Shannon diversity) were assessed using ANOVA. Beta diversity (Bray-Curtis dissimilarity) was analysed using multivariate analysis of community separation (PERMANOVA, using the Adonis function), with pre-exposure and adult exposure as fixed factors and 99,999 permutations. Statistical differences in ASV abundance were evaluated using DESeq2 [11]. Low abundance ASVs were independently filtered within DESeq2, and default settings were applied for outlier detection and dispersion moderation. ASVs were considered significantly differentially abundant at an adjusted P value $<0.05$.
